# Peripheral Blood Nascent RNA Sequencing Captures Ancestry-Linked Differential Gene Expression

**DOI:** 10.64898/2026.01.15.699738

**Authors:** Aileen Kimm, Emily Gini Uh, Hojoong Kwak

## Abstract

Genetic ancestry reflects inherited population structure that can influence gene regulation and disease susceptibility, yet biomedical studies continue to rely on self-identified race as a proxy for genetic variation. We developed a streamlined framework to infer local and global genetic ancestry directly from nascent RNA sequencing data and tested whether ancestry-based stratification improves detection of transcriptional differences compared with race. Peripheral blood samples from 50 donors were profiled using Peripheral Blood Chromatin Run-On sequencing (pChRO). Ancestry-informative single nucleotide polymorphisms were identified from aligned reads and compared with 1000 Genomes reference populations representing populations in the U.S. using a Bayesian model to infer ancestry locally. Global ancestry proportions were assigned by unsupervised clustering methods. Differential gene expression was analyzed by controlling for age and sex, and the number of true differentially expressed genes (DEGs) was estimated using a q-value framework. Ancestry inference from sparse nascent transcription data recovered heterogeneous local ancestry patterns and separated individuals into two major ancestry clusters, revealing multiple discordances with self-identified race. Grouping by genetically inferred ancestry increased power to detect transcriptional differences relative to to self-identified race, including genes involved in dermatologic and neurodegenerative pathways. These results demonstrate that nascent RNA sequencing enables ancestry inference and that ancestry-based stratification captures transcriptional variation not resolved by race, supporting its integration into population scale functional genomics study.

## INTRODUCTION

Genetic ancestry, identified from genome-wide patterns of inherited variations, captures biologically meaningful population structure that can influence gene regulation, disease susceptibility, and treatment response. As omics-based medicine moves towards personalized approaches, integrating ancestry-informed biological differences becomes increasingly important. Omics studies span a broad range of molecular layers, from genetic and epigenetic variation across individuals and populations to downstream consequences of genetic and environmental factors, including proteomic or metabolomic profiles (1). Integrative multi-omics approaches provide opportunities to incorporate genetic ancestry alongside these molecular features to enhance our understanding of disease mechanisms and therapeutic strategies (2).

Historical use of race in biomedical research and clinical practice often faced fundamental limitations, as race is a social construct shaped by political and cultural context rather than stable biological categories (3,4). Earlier racial classifications often relied on inconsistent or exclusionary criteria contributing to the systematic marginalization of minority groups, notably skin color (5,6). The effects of racial discrimination often persist through socioeconomic factors, such as income level and quality of living conditions, and have been correlated with poorer health outcomes for minority groups (7). While efforts to combat health disparities through more personalized medicine has led to the inclusion of race in clinical equations, such for as estimated glomerular filtration rate (GFR) or lung function, these methods are now being reevaluated with support from philanthropies such as the Doris Duke Foundation (8,9). These reassessments reflect growing recognition that race does not reliably represent underlying genomic variation or biological determinants of health. Although self-identified race and ethnicity provide valuable information about social and lived experience, they remain imprecise proxies for inherited genetic structure.

This ambiguity underscores persistent challenges in multi-omics research, which already faces limitations in data standardization and notable under-representation of non-European populations, despite initiatives such as H3Africa and GenomeAsia 100K (10,11). As the field seeks more accurate and equitable analytical frameworks, genetic ancestry offers a biologically informative framework for capturing population structure relevant to gene regulation and disease. Used carefully, genetic ancestry may provide a more objective yet sociologically informed alternative to race for integrating human diversity into omics analyses and advancing personalized medicine.

As multi-omics methodologies expand, nascent RNA–based approaches offer a practical and unexpectedly powerful means of genetic ancestry inference directly from patient samples. Traditionally, genetic ancestry estimation relies on high-quality genomic DNA obtained from peripheral blood, followed by genotyping Ancestry-Informative SNP Markers (AIMs) and comparison to reference populations using tools such as ADMIXTURE or STRUCTURE (12). These pipelines are optimized for DNA-based datasets typically generated explicitly for ancestry inference, limiting their integration with routine sequencing workflows designed for other purposes. In contrast, datasets from nascent RNA profiling technologies such as Precision Run-On Sequencing (PRO-seq) capture a substantially larger fraction of the genome than conventional expression assays, such as mRNA-seq, due in part to their extensive coverage of intronic regions (13). This broader genomic coverage increases the representation of AIMs, thereby enabling genetic ancestry inference from nascent RNA sequencing data.

Since PRO-seq and pChRO can be performed directly on whole blood or other minimally processed biospecimens, they might provide a practical and accessible way to obtain genome-wide transcriptional data that inherently contains the same inherited polymorphisms needed to identify ancestry-informative markers. As a result, nascent RNA datasets—originally produced to study real-time transcriptional regulation—can also support both local and global genetic ancestry inference without additional DNA-specific sample preparation or genotyping. Leveraging nascent RNA for ancestry inference broadens access to genetic structure information and integrates seamlessly within multi-omics frameworks, connecting population-specific regulatory variation with transcriptional dynamics from the same dataset.

In this study, we present a streamlined approach for inferring genetic ancestry using nascent RNA. We analyzed fifty frozen peripheral blood donor samples with Precision Run-On Sequencing (PRO-seq) to identify AIMs and map them to reference populations in Haplotype Map (HapMap). As a proof of concept, and to extend prior studies examining population-associated genetic variations and expression differences (14,15), we further assess the differential expression of genes implicated in human diseases.

## METHODS

### Study participants and data pChRO data processing

Deidentified human peripheral blood samples (n = 50) were purchased from Innovative Research (IR) Inc. Samples were provided with limited metadata, including self-identified race, sex, and age. The specimens were collected and fully de-identified under the compliance of institutional review board (IRB)–approved protocols by IR Inc., and were considered exempt from additional IRB reviews.

Approximately 2-3 mL from each donor sample was used for nascent RNA profiling with Peripheral Blood Chromatin Run-On sequencing (pChRO), a modification of the Precision Run-On sequencing (PRO-seq) protocol optimized for peripheral leukocytes. Briefly, 3-4 ml of frozen whole blood was thawed on ice and immediately lysed in 20 ml of NUN buffer (0.3 M NaCl, 1 M urea, 1% NP-40, 20 mM HEPES pH 7.5, 7.5 mM MgCl_2_, 0.2 mM EDTA, 1× protease inhibitor cocktail, 1 mM DTT, and 4 U/ml RNase inhibitor) with vortexing. Chromatin was pelleted by centrifugation at 15,000 × g for 20 min at 4 °C and resuspended in Wash Buffer (50 mM Tris-HCl, pH 7.5). After brief centrifugation, the pellet was washed once in Buffer D (50 mM Tris-HCl, pH 8.0, 25% glycerol, 5 mM magnesium acetate, 0.1 mM EDTA, 5 mM DTT). The chromatin pellet was then homogenized by brief sonication (10 x 1 s on/ 1 s off pulses, tip sonicator) in 50 μl of Buffer D and immediately used for nuclear run-on reactions, and subsequent PRO-seq libraries were generated as described previously (16,17).

Each library was sequenced on an Illumina platform, producing 75 bp single-end FASTQ files. Reads were aligned to the human reference genome (hg38), and the alignments were stored in binary alignment and map (BAM) format. Single nucleotide polymorphisms (SNPs) at AIMs were identified by comparison to the reference genome and classified as either *reference* (matching to hg38) or *alternative* (non-matching). SNP data from all 50 donor samples were aggregated into non-overlapping genomic windows of 2 megabases (Mb) and served as input for downstream ancestry inference analysis.

### Reference populations and local ancestry inference

Population reference haplotype maps were obtained from the 1000 Genomes Project (18). Variant call format (VCF) files corresponding to Mexican ancestry in Los Angeles (MXL), African-American ancestry in the Southwest US (ASW), and Utah residents with Northern and Western European ancestry (CEPH) were downloaded and used as reference populations. For each variant SNPs, allele frequencies were computed to estimate the probability of observing an alternative allele in each reference population relative to hg38, which were used in the Bayesian modeling in subsequent steps AIMs were filtered using two criteria: minor allele frequency (MAF) 5% in at least one population and inter-population allele frequency difference > 25% between at least one pair of populations (19). A total of 3,331,306 AIM SNPs and reference probabilities for each population were identified.

Filtered AIMs were compared to sample SNPs from the pChRO data within each non-overlapping genomic window. Only ancestry-informative markers (AIMs) with at least one pChRO read covering the variant position, sufficient to determine the reference or alternative allele, were included for each individual. For each window, the Bayesian posterior probability of ancestry assignment to the MXL, ASW, or CEPH populations were calculated based on the likelihood of observed alleles given the population-specific allele frequency profiles. Each window was assigned to the ancestry group with the highest posterior probability, and window-level probabilities were summed to infer local genetic ancestry across the genome. Genome-wide ancestry proportions were estimated by averaging ancestry probability distributions across all genomic windows.

### Cluster analysis and significance tests

Concordance between self-identified race and genetically inferred ancestry was first assessed using k-means clustering. Two optimal clusters were identified (20), along with mismatches and outlier samples. To validate and visualize clustering structures, t-distributed stochastic neighbor embedding (t-SNE) was performed.

### Differential Gene Expression Analysis

Gene expression across ancestry groups was analyzed using DESeq2, with sex and age included as covariates to control for potential confounding effects. Samples with mismatched ancestry (based on clustering) were reassigned to their genetically inferred ancestry group to enable comparison between models using self-identified race and genetically inferred ancestry. Raw p-value histogram plots were generated to visualize the distribution of *p*-values across all genes, providing an assessment of statistical power contrasting ancestry-based clustering with race-based grouping.

The true number of differentially expressed genes (DEGs) was estimated from the distribution of raw p-values using the method of Storey and colleagues (15). Briefly, p-values were filtered to remove missing values, and the proportion of null hypotheses (π_0_) was estimated across a range of tuning parameters (λ=0.05–0.95) based on the fraction of p-values exceeding each λ. The resulting π_0_(λ) estimates were smoothed using spline regression, and the final π_0_ value was obtained by evaluating the fitted curve at the maximum λ. The number of true DEGs was then calculated as m(1−π_0_), where m is the total number of tests. In addition, false discovery rate–adjusted q-values were computed using Storey’s direct FDR approach to enable gene-level significance assessment.

Functional enrichment analysis of differentially expressed genes (FDR < 0.1) was performed using the PANTHER Gene Ontology database (pantherdb.org).

## RESULTS

### Local and global ancestry inference from nascent transcription data

We inferred genetic ancestry for 50 individuals using sparse nascent transcription data and a Bayesian framework with reference populations from the 1000 Genomes Project (1KGP) dataset, including African Ancestry in Southwest United States (ASW), Utah Residents with Northern and Western European Ancestry (CEU), and Mexican Ancestry in Los Angeles (MXL). Local ancestry inference results for representative examples (individuals 12, 20, 21, and 31) are shown (**Figure 1A**).

**Figure 1.**
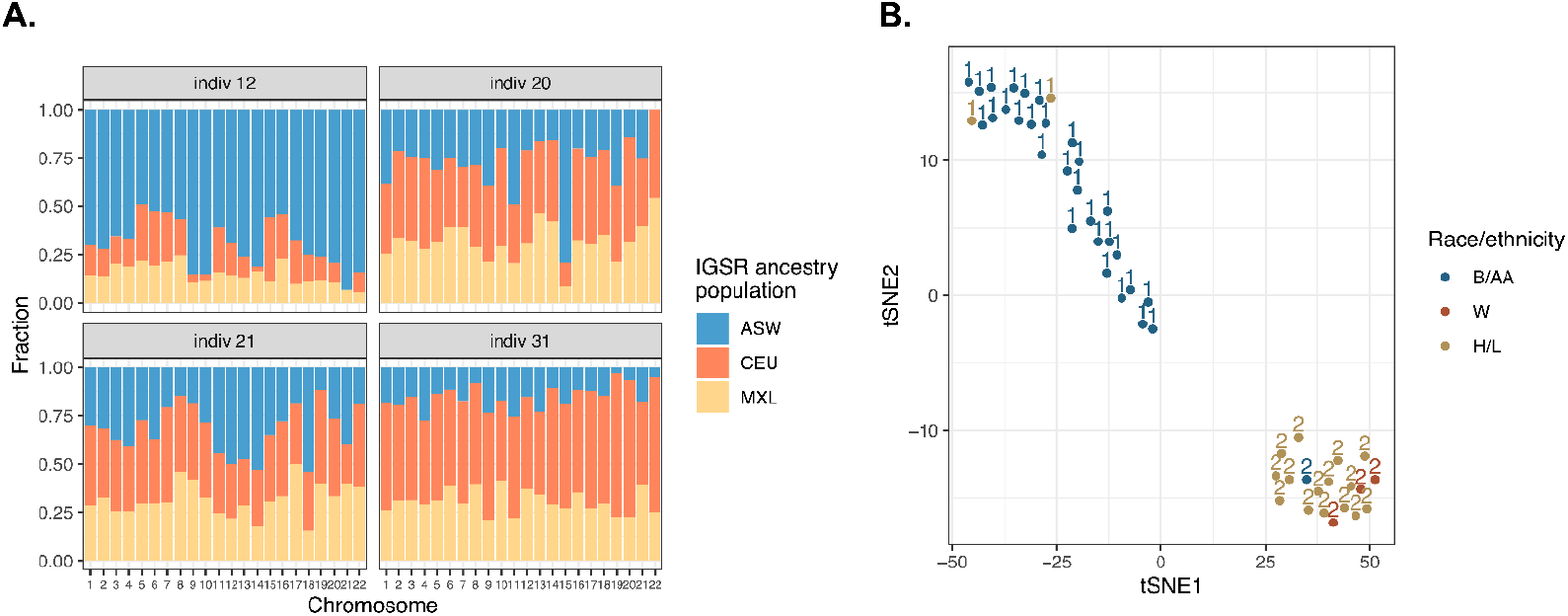
Local and global ancestry inference. **A.** Local ancestry inference in select individuals. Color bars indicate proportions of ancestry population with highest Bayesian probability for 2 megabase block arrays in each chromosome. IGSR: ASW (African Ancestry in Southwest United States), CEU (Utah Residents with Northern and Western European Ancestry), and Mexican Ancestry in Los Angeles (MXL). **B**. Global ancestry clustering of individuals based on t-SNE of genome-wide ancestry proportions. Points are colored by self-reported race/ethnicity: B/AA (Black/African American), W (White), and H/L (Hispanic/Latino). Distinct clusters reflect differences in global ancestry composition.

Local ancestry profiles revealed heterogeneous ancestry contributions across chromosomes, consistent with admixture patterns captured by the reference populations. For example, individual 12 exhibited predominantly ASW ancestry across the genome, with smaller contributions from CEU and MXL, consistent with their self-identification as African American (AA). Global ancestry estimates for this individual were 0.72 ASW, 0.13 CEU, and 0.15 MXL. The sums of ancestral proportions approximated to the expected value of 1, indicating stable Bayesian inference and appropriate representation by the reference populations.

To assess global ancestry structure across individuals, we performed t-distributed stochastic neighbor embedding (t-SNE) using genome-wide ancestry proportions (**Figure 1B**). This analysis separated samples into two major clusters. Cluster 1 was characterized by predominantly ASW ancestry, whereas Cluster 2 showed substantial MXL ancestry with additional CEU contributions. These clusters reflect differences in global ancestry composition rather than self-reported race or ethnicity.

Notably, discrepancies were observed between self-identified race/ethnicity and genetically inferred ancestry in some individuals. For example, individual 9, who self-identified as Hispanic/Latino, clustered with the ASW-dominant group (Cluster 1) in both technical replicates. Conversely, individual 10, who self-identified as African American, clustered with the MXL/CEU-enriched group (Cluster 2). These cases highlight the limitations of self-reported race as a proxy for genetic ancestry and underscore the ability of nascent RNA–derived data to recover ancestry-associated genetic structures.

### Ancestry-based stratification enhances detection of transcriptional differences

Differential gene expression (DGE) analysis revealed substantially greater statistical power when individuals were grouped by inferred genetic ancestry rather than by self-identified race (**Figure 2**). Because self-reported race does not directly encode shared genomic structure, it may introduce heterogeneity in ancestry-related analysis. In contrast, ancestry inference leverages population-specific allele distributions and therefore more directly captures genetic variation relevant to transcriptional regulation. As a result, race-based models are more susceptible to misclassification, particularly in admixed individuals or in genomic regions with heterogeneous or rapidly shifting local ancestry, reducing sensitivity for detecting true expression differences.

**Figure 2.**
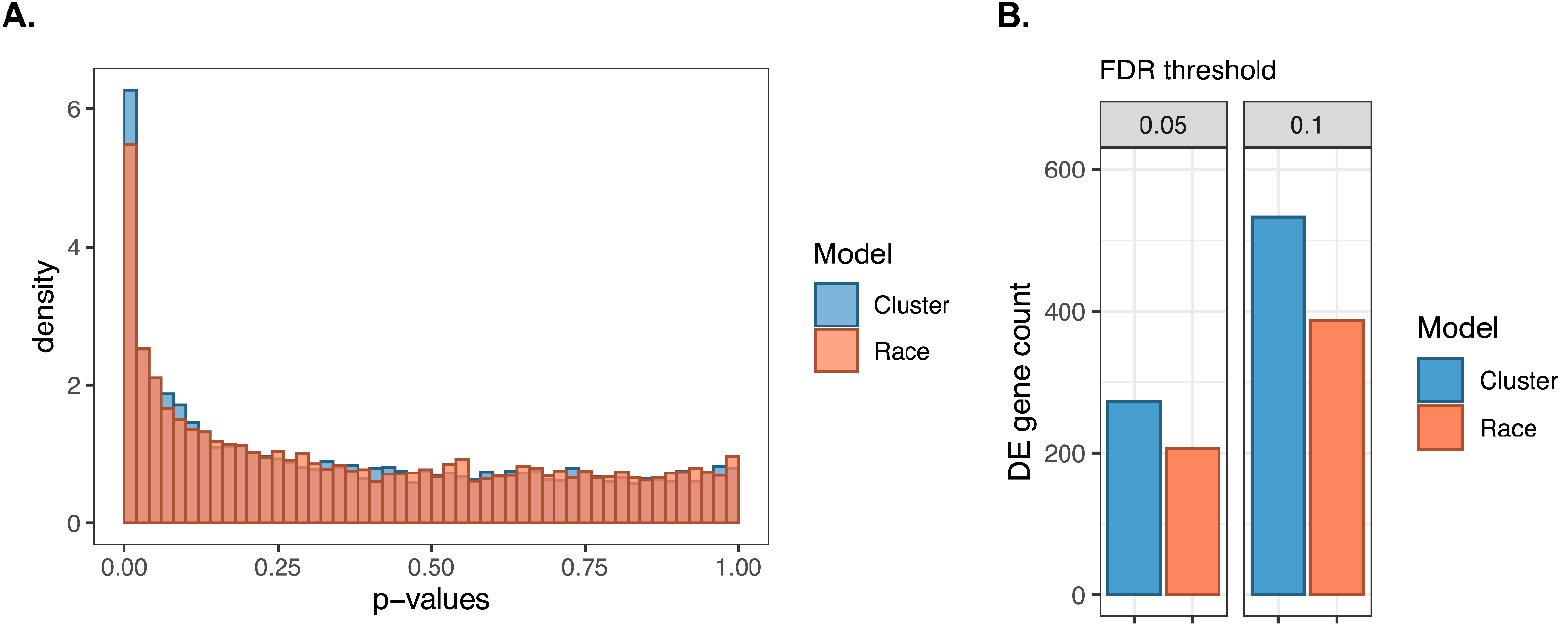
Differential gene expression between genetic ancestry clusters. **A.** Raw p-value histogram comparing differential gene expression based on genetic ancestry cluster (cluster) or self-identified race (race). **B**. Number of differentially expressed genes based on cluster or race. (p-values using McNemmer test, p < 1.22 x 10^-6^, p < 1.1 x 10^-15^ respectively for FDR thresholds 0.05 and 0.1)

This difference in statistical power is evaluated using distribution of raw p-values (**Figure 2A**). The ancestry-based cluster model showed a stronger enrichment of small p-values compared with the race-based model, consistent with increased signal relative to noise. To estimate the number of true differentially expressed genes (DEG), we applied a q-value-based approach that infers the proportion of null hypotheses from the p-value distribution. This q-value–based method estimates the overall number of true differentially expressed genes by inferring the proportion of null hypotheses from the p-value distribution, while individual genes were classified as differentially expressed using conventional false discovery rate (FDR)-adjusted p-value thresholds. Using these methods, the cluster model estimated 1,637 true DEGS (26%), whereas the race model estimated 1,438 true DEGs (23%)(21,22).

Consistent with these findings, the total number of DEGs passing conventional FDR thresholds was significantly higher for ancestry-based clustering than for race-based grouping across both FDR < 0.05 and FDR < 0.1 (**Figure 2B**). The improvement in DEG recovery using ancestry-based clustering was therefore statistically significant (Fisher’s exact test, p = 3.90 × 10^−5^), indicating that inferred genetic ancestry more effectively stratifies individuals according to biologically meaningful transcriptional differences than self-reported race.

### Survey of ancestry-associated transcriptional variation

To characterize transcriptional differences associated with genetic ancestry, we surveyed differentially expressed genes between ancestry-defined clusters (**Figure 3A**). Among 6,248 genes expressed above 1 tag per million mapped reads (TPM) across 50 individuals (background gene set), 519 (8.3%) were differentially expressed at an adjusted p-value < 0.1. Notably, these ancestry-associated DEGs are enriched with disease associated gene sets. For example, of the 57 genes annotated as skin-related in our background gene set, 11 (19.3%) were differentially expressed, indicating a significant enrichment of dermatologically relevant genes (p=0.019, Fisher exact test).

**Figure 3.**
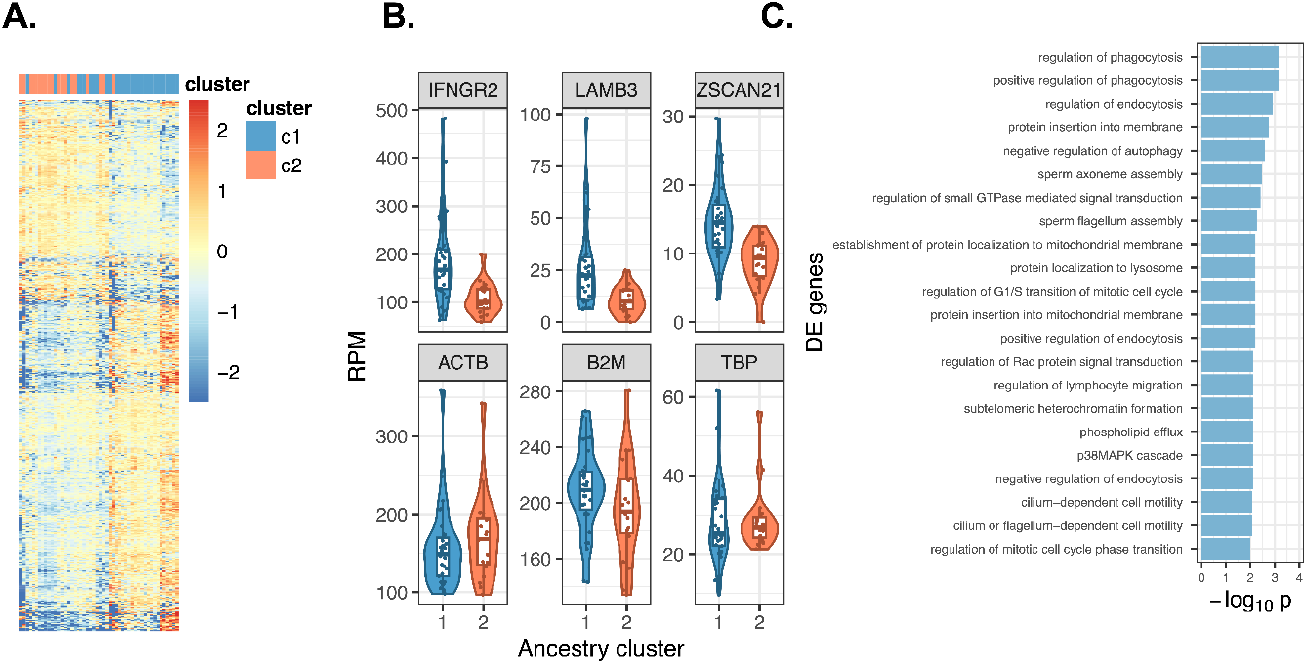
Differential gene expression and pathway enrichment between genetic ancestry clusters. **A.**Heatmap of normalized gene expression (row-scaled Z-scores) for genes differentially expressed between ancestry clusters c1 and c2 at a false discovery rate (FDR) < 0.1, identifying 519 differentially expressed genes (DEGs). **B**. Violin plots showing expression distributions (RPM) for representative DEGs across clusters, including IFNGR2, LAMB3, and ZSCAN21. Housekeeping genes ACTB, B2M, and TBP are not significantly different between clusters (ACTB FC = 0.12, p = 0.319; B2M FC = −0.11, p = 0.149; TBP FC = −0.34, p = 0.265). **C**. Gene ontology enrichment analysis of DEGs

Expression levels of canonical housekeeping genes – *ACTB, B2M*, and *TBP* – were examined as internal controls (**Figure 3B**). These genes encode β-actin (a core cytoskeletal component), β2-microglobulin (a structural element of MHC class I molecules), and TATA-binding protein (a general transcription factor), respectively, are typically stable across biological conditions. Consistent with expectation, none of these showed significant differences in gene expression between the two ancestry clusters, supporting the specificity of the observed differential expression patterns..

In contrast, DEGs showed pronounced ancestry-associated expression differences. Among those were genes with known relevance to dermatological and neurodegenerative conditions (**Figure 3B**). Two skin disease-related genes, *IFNGR2* and *LAMB3* (23,24), were more highly expressed in cluster 1 compared to cluster 2 (p=3.06 × 10^−6^ and 2.14 × 10^−6^; fold changes of approximately 1.7 and 2.6, respectively). These genes encode proteins involved in immune signaling and epidermal structural integrity. Similarly *ZSCAN21*, associated with Parkinson’s disease (25,26), showed higher expression in cluster 1(p = 8.27 × 10^−7^; approximately 1.7 fold-increase). Gene ontology analysis of differentially expressed genes enrichment for biological processes related to phagocytosis, endocytosis, membrane trafficking, and autophagy (**Figure 3C**). Together, these results indicate that ancestry-informed stratification captures biologically meaningful transcriptional variation, including pathways relevant to skin biology and cellular homeostasis.

## DISCUSSION

In developing our method, our primary goal was to investigate the ability to utilize nascent RNA to infer genetic ancestry data for the purpose of verifying the strength of using genetic ancestry rather than race in categorizing patient samples. Despite the sparse data provided by PRO-seq, we were able to determine both local and global ancestry for each sample and compare these with the self-identified race of the individual. Through tSNE clustering, we identified discrepancies between race and ancestry categorization for several samples (**Figure 1B**). Additional analysis validated the greater accuracy of grouping samples by ancestry, as seen by comparing statistically significant differential gene expression (**Figure 2B**). This finding was reflected in examining individual genes as well. As expected for a control group, the gene expression for housekeeping genes remained similar across samples regardless of grouping by ancestry or race. On the other hand, expression levels were stronger for genes known to be associated with specific disorders (e.g. LAMB3, IFNGR2, ZSCAN21) (**Figure 3B**).

The higher expression of genes that are specific to individual clusters can provide reasonable evidence for genetic ancestry and its potential role in disease prevalence. If gene expression patterns reflect underlying biological pathways implicated in disease, ancestry-associated regulatory differences may lead to further investigation in disease-focused cohorts. The observed expressions of IFNGR2, LAMB3, and ZSCAN21 largely align with epidemiologic observations of population-level differences in disease prevalence (23–26), suggesting that contemporary populations may serve as imperfect but informative proxies for ancestral populations. However, this approach is limited by the use of self-identified race as a surrogate for genetic ancestry, which does not fully capture population heterogeneity or admixture.

In Figure 3B, IFNGR2 expression was significantly elevated in cluster 1, consistent with the higher prevalence of immune-mediated dermatoses such as lichen planus reported in African Americans (27). IFNGR2, which forms part of the interferon-γ receptor complex, participates in host defense and inflammatory pathways and is implicated in lichen planus (28) and psoriasis (29). LAMB3 expression was also significantly increased in cluster 1, suggesting potential ancestry-associated contribution to epidermal or basement membrane-related pathways, although direct links to disease prevalence remain to be established. LAMB3, which encodes the β3 subunit of laminin-332, is a critical component of the epidermal basement membrane; pathogenic variants lead to junctional epidermolysis bullosa (30), and its overexpression has been associated with increased risk of cutaneous squamous cell carcinoma (23). Together, the high proportion of differentially expressed genes across clusters is consistent with ancestry-associated differences in immune and skin-related transcriptional pathways, underscoring the importance of genetic ancestry in understanding disease susceptibility.

ZSCAN21 encodes a zinc finger and SCAN domain-containing transcription factor that regulates neuronal gene expression and has been associated with susceptibility to Parkinson’s disease through modulation of dopaminergic signaling pathways (25, 26). Expression of ZSCAN21 also differed between the two clusters, further illustrating how genetically inferred ancestry can reveal subtle but biologically meaningful transcriptional variation across disease-relevant gene sets. Our finding is supported by a broader genome-wide association analysis of Parkinson’s disease, which also detected significant variation between ancestral groups for numerous related genes (31).

### Race and Genetic Ancestry

Our results align with the growing understanding within literature that genetic ancestry and race are not interchangeable (32). While there are set protocols for ancestry inference (e.g. analyzing AIMs within a sample genome in comparison to reference genomes), race is deeply intertwined with an individual’s perspective of their identity and culture. Similarly, the perception of race is also influenced by the sociopolitical climate, as seen by consistent updates in the US census standards for race/ethnicity data collection. In 2024, “Middle Eastern and North African” was introduced as a new category, rather than its previous integration into the “White” category (33). Although race correlates with important socioeconomic factors (7), it may be limited as a variable in studies capturing inherited genetic variations.

In our study, four individuals (**Figure 1B**) seem to represent this distinction, as their self-identified race does not correlate with the expected ancestry group. The nuances of race are further highlighted with inspection of the distribution of ancestral groups for each individual (**Figure 1A**). One notable example was individual 31, who self-identified as African-American but was found to have similar percentages of CEU and MXL ancestry, alluding to admixture, or the interbreeding of two previously separate ancestral groups (34). For this individual, identifying as African-American may be a result of sociological factors (e.g. culture and environment) rather than ancestry.

Admixture and difficulties with defining race remain as persisting reasons for the steady transition away from race-derived clinical equations (8). Omics-based medicine strives to utilize a more objective method towards the same principles of precision medicine. In examining the validity of clinical equations, some studies have argued for the use of alternative measurements (e.g. cystatin C values rather than race as a risk factor for kidney disease (35)), while others have found that genetic ancestry can be a more accurate, alternative patient factor (e.g. when determining lung function values (36)).

### Limitations and Future Directions

The socially constructed nature of race, and its variable interpretation across individuals and contexts, presents practical challenges in obtaining reliable and consistent racial metadata across human samples. In this study, self-identified race collected by third-party vendors was provided alongside de-identified specimens, with limited access to specific survey instructions and methods to collect these data on the researcher’s side. Prior studies have shown that survey wording can substantially influence responses to questions about race (37), and participants’ interpretation of race categories may not align with the survey’s intentions. For example, individuals identifying with multiple racial or ethnic categories (e.g. African American, Hispanic/Latino, and White) may have selected solely one category due to ambiguities in survey instructions and the definition of each category. Furthermore, as each sample was de-identified, labeling errors introduced during sample collection or annotation cannot be excluded as a source of race-ancestry discrepancies.

We also recognize that genetic ancestry inference itself is subject to systematic limitations. Reference panels used in ancestry assignment, such as the Haplotype Map, often rely on labels generated from geographic sampling locations (e.g. “Peruvians in Lima, Peru” (18)) rather than clearly defined or historically contextualized population groups. The use of broad continental ancestry has been criticized for the risk of conflation with socially constructed racial categories and for oversimplifying human demographic histories (38). In this sense, incomplete knowledge of historical migration patterns and admixture events may limit the resolution and interpretability of ancestral assignments (39). This issue is further compounded by under-representation of non-European populations in widely used genetic databases, which can reduce inference accuracy and generalizability for individuals with diverse ancestral backgrounds (11).

Importantly, the goal of our approach is not to promote genetic ancestry as a direct replacement for race, but rather position genetic ancestry as one informative component within a broader precision medicine framework. By providing accessible and scalable methods for estimating genetic ancestry, we aim to support more nuanced study designs that reduce reliance on race as a biological proxy and instead evaluate whether genetically informed measures may better capture relevant sources of biological variations in their studies.

